# Bacterial Microbiome and Host Inflammatory Gene Expression in Foreskin Tissue

**DOI:** 10.1101/2022.08.29.505718

**Authors:** Brandon S. Maust, Stefan Petkov, Carolina Herrera, Colin Feng, Bryan P. Brown, Limakatso Lebina, Daniel Opoka, Andrew Ssemata, Natasha Pillay, Jennifer Serwanga, Portia Seatlholo, Patricia Namubiru, Geoffrey Odoch, Susan Mugaba, Thabiso Seiphetlo, Clive M. Gray, Pontiano Kaleebu, Emily L. Webb, Neil Martinson, Francesca Chiodi, Julie Fox, Heather B. Jaspan, the CHAPS team

## Abstract

As part of the CHAPS randomized clinical trial, we sequenced a segment of the bacterial 16S rRNA gene from foreskin tissue of 144 adolescents from South Africa and Uganda collected during surgical penile circumcision after receipt of 1 to 2 doses of placebo, emtricitabine with tenofovir disoproxil fumarate, or emtricitabine with tenofovir alafenamide. We found a large proportion of *Corynebacterium* in addition to other anaerobic species. *Cutibacterium acnes* was more abundant among participants from South Africa than Uganda, though this made no difference in surgical recovery. We did not find a difference in bacterial populations by treatment received nor bacterial taxa that were differentially abundant between participants who received placebo versus active drug. Using RNAseq libraries from foreskin tissue of the same participants, we found negative correlations between the relative abundance of bacterial taxa and the expression of genes downstream of the innate response to bacteria and regulation of the inflammatory response. When participants were divided into clusters based on bacterial community composition, two main clusters emerged which were distinguished by high and low bacterial diversity. Random forest classification showed higher expression of *NFATC3* and *SELENOS* and lower expression of *STAP1* and *NLRP6* in the higher diversity group compared to the lower. Our results show no difference in the tissue microbiome of the foreskin with short-course PrEP but that bacterial taxa were largely inversely correlated with gene expression, consistent with non-inflammatory colonization.

**Author Summary:** We investigated the bacterial community of the foreskin of the penis. Previous studies found increased inflammation with certain anaerobic bacteria from swabs taken under the foreskin, but we found that higher relative abundances of the bacteria were correlated with lower expression of inflammatory genes. We did not find different bacteria in participants who received medicine to prevent HIV. Understanding the relationship between bacteria and inflammation in the penis will help us to understand how interventions like penile circumcision reduce the risk of acquiring sexually transmitted infections such as HIV.

## Introduction

HIV remains a significant global health challenge despite substantial clinical, public health, and basic science research efforts. While condom use (1, 2), pre-exposure prophylaxis (PrEP) (3), and medical penile circumcision (4) are effective at reducing HIV incidence in men, the contribution of the penile microbiome to these mechanisms has not been fully explored. Previous work has shown a predominance of anaerobic species in the microbiota of swabs taken from the coronal sulcus or urethra (5-15) and have reported associations between species such as *Prevotella* and increased mucosal inflammation, HIV target cell density and risk of HIV acquisition (8, 16). Following circumcision, the surface microbiota shifts to be dominated by more aerobic species as found on other skin surfaces (10, 17, 18). Thus far, no data exist as to the microbiota of the foreskin itself, and its relation to tissue inflammation.

Antiretrovirals (ARVs) are used for both treatment and prevention of HIV, and limited data have shown a complex relationship with bacteria. When applied as topical vaginal pre-exposure prophylaxis, *L. crispatus* was shown to endocytose Tenofovir (TFV) then either actively metabolize or release it back into the environment (19). Similarly, *Gardnerella vaginalis* and other anaerobes have been shown to metabolize TFV (20) or block its entry into cells by secretion of adenine (19). Antiretrovirals may also theoretically alter bacteriophage populations which can dramatically reshape the bacterial component of the microbiome which they infect (21). At the rectal mucosa, small studies have investigated the effects of oral emtricitabine (FTC) with tenofovir disoproxil fumarate (TDF) on the bacterial microbiome and innate inflammatory pathways in men who have sex with men (MSM) and transwomen (22-25) but with varying results.

Within the CHAPS clinical trial, young men were randomized to 1 to 2 doses of placebo, emtricitabine with tenofovir disoproxil fumarate, or emtricitabine with tenofovir alafenamide, prior to medical penile circumcision. Foreskin tissue was collected and subject to both 16S rRNA sequencing and RNASeq. to characterize the bacterial microbiome and inflammatory gene expression. We hypothesized that short courses of PrEP, as utilized in a dose-finding trial, would not result in significant changes to the bacterial microbiome of the foreskin, but that there would be a relationship between the microbiota and inflammatory gene expression.

## Materials and Methods

### Cohort and Specimen Collection

The Combined HIV Adolescent PrEP and Prevention Study (CHAPS) was a randomized controlled trial that enrolled 144 men living without HIV aged 13-24 years between 2019 and 2021 from the Chris Hani Baragwanath Academic Hospital in Soweto, South Africa (n=72) and the Entebbe Regional Referral Hospital in Entebbe, Uganda (n=72). Inclusion criteria were male sex at birth, hemoglobin > 9 g /dL, weight > 35 kg, two successive negative rapid HIV antibody tests, and clinical eligibility for surgical circumcision (26). Exclusion criteria were conditions precluding circumcision or receipt of the study medications. Participants were randomized to placebo versus FTC with either TDF or tenofovir alafenamide (TAF) for 1-2 days prior to surgical penile circumcision to investigate ARV dosing for on-demand PrEP.

All participants underwent a physical exam at study entry and completed survey instruments including sexual history at the randomization visit. At the circumcision visit, they provided midstream urine for *Chlamydia trachomatis* (CT) and *Neisseria gonorrhea* (GC) testing via nucleic acid amplification testing (NAAT) prior to surgery. If an asymptomatic sexually transmitted infection was diagnosed, antibiotic treatment was prescribed at the post-operative visit. Penile circumcision was performed using the dorsal slit method and the removed prepuce was placed immediately in cold Dulbecco’s Modified Eagle Medium and shipped on ice within 1 hour (median 30 minutes) to the local laboratories in Uganda (Medical Research Council/Uganda Virus Research Institute) and South Africa (Perinatal HIV Research Unit in Johannesburg) for processing. Smaller, 5–7 mm^2^-sized sections were stored dry at −80° C until the samples were transported on dry ice to the Seattle Children’s Research Institute, U.S.A for microbiome studies and to the Karolinska Institutet, Sweden for transcriptome analyses.

### Ethics and Human Subjects

Ethical clearance to conduct the CHAPS trial was obtained from the South African Health Products Regulatory Authority (20181004), the Uganda Virus Research Institute research and ethics committee (GC /127 /18 /12 /680), Uganda National Council of Science and Technology (HS 2534), Uganda National Drug Authority (618 /NDA /DPS /09 /2019), and the London School of Hygiene and Tropical Medicine research ethics committee (Ref:17403). Informed written consent was collected from all participants. The Swedish Ethics Review Authority approved the transcriptome studies of the collected specimens at the Karolinska Institutet (2020-00941). The ethics approval for the microbiome analysis was granted by the Seattle Children’s Institutional Review Board (STUDY00003430).

### Specimen Processing

#### 16S rRNA Analysis

At the time of analysis, vials were thawed and approximately 25 mg of tissue was dissected and processed by a customized Qiagen PowerSoil Pro protocol for extraction of DNA using a QIAcube instrument, available at: https://dx.doi.org/10.17504/protocols.io.4r3l2774jg1y/v1. A negative extraction control consisting of solution CD1 without specimen was also included. The specimens from each collection site were extracted on single plates. The resulting total DNA was diluted 1:4 to reduce PCR inhibition.

The *16S* rRNA gene V3-V4 region was amplified using 319F / 806R universal primers for 20 cycles of PCR as previously described (27) for each specimen along with a negative PCR control reaction consisting of mastermix without DNA template for each replicate and evenly and staggered genomic DNA from mock bacterial libraries (BEI Resources) as positive sequencing controls. The amplified products were purified using Agencourt AMPure XP beads (Beckman Coulter) and submitted to an additional 10 rounds of PCR with indexing primers (Illumina). The resulting libraries were pooled by volume with specimens at 100x the positive controls. The resulting library comprising all participants was purified using a MinElute PCR purification column (Qiagen), followed by the QiaQuick gel extraction kit (Qiagen). The cleaned library was quantitated using qPCR (NEBNext Library Quant Kit for Illumina), then pooled with PhiX, denatured, and loaded onto a MiSeq instrument (Illumina) with a v3 2×300 flow cell following the manufacturer’s protocol.

Sequences were de-multiplexed using Illumina’s BaseSpace workflow. Primers and adapters were removed by cutadapt 2.7 (28). Sequences were further trimmed for quality, then filtered and merged using dada2 1.22.0 (29) to generate amplicon sequence variants (ASVs). Taxa were annotated using the Silva 138.1 database (30) with additional genital-associated species (31) using a 100% nucleotide identity threshold. The phyloseq 1.40.0 (32) and vegan 2.6-2 (33) R packages were used to manipulate ASV tables and calculate diversity measures. ASV sequences were aligned using ssu-align 0.1.1 (34), and a maximum likelihood phylogeny was generated using PhyML 3.3.20220408 (35) with a GTR substitution model. Contaminating sequences were identified by their presence in negative controls for the extraction and PCR amplification or mock community using decontam 1.16.0 (36) and microfiltR (37). After decontamination, specimens with fewer than 25-fold as many reads than extraction and PCR controls were excluded. For differential abundance analysis, decontaminated ASVs were filtered with prevalence >= 10% and relative abundance threshold of 1×10 ^-4^ before combining counts for all ASVs classified as the same species. ALDEx2 1.28.1 (38), ANCOM-BC 1.6.2 (39), and DESeq2 1.36.0 (40) (using the poscounts factors estimation) were used for differential abundance testing to overcome the documented limited power and accuracy of these tools when used individually on 16S data sets which contain a high proportion of zero counts (41, 42).

#### RNAseq of Foreskin Tissue

Foreskin samples were disrupted and homogenized using a Tissuelyzer (Qiagen) and total RNA isolated using the RNeasy Kit (Qiagen) according to manufacturer’s instructions. RNA was subjected to quality control with Agilent Bioanalyzer (Agilent). To construct libraries suitable for Illumina sequencing, the Illumina stranded mRNA prep ligation sample preparation protocol was used with starting concentration of 200 ng total RNA. The protocol includes mRNA isolation, cDNA synthesis, ligation of adapters and amplification of indexed libraries. The yield and quality of the amplified libraries were analysed using Qubit by (Thermo Fisher) and the Agilent Tapestation (Agilent). The indexed cDNA libraries were normalized and combined, and the pools were sequenced on the Illumina Novaseq 6000 S4 flowcell to generate 150 bp paired-end reads.

Sample demultiplexing was performed using bcl2fastq 2.20.0 (Illumina), and quality and adapter trimming of reads was performed using Cutadapt 2.8 (28). Sample quality was assessed using FastQC 0.11.8 (Babraham Bioinformatics) and MultiQC 1.7 (43). Reads were aligned to the Ensembl GRCh38 reference genome using STAR 2.6.1d (44). Counts for each gene were obtained using featureCounts 1.5.1 (45).

The Gene Ontology (GO) term “inflammatory response” (GO:0006954) selected 860 putative inflammatory genes which were filtered to only those with at least two copies detected in at least 90% of specimens. RNA read counts were normalized then transformed by centered log ratio (CLR). The 16S ASVs were filtered and combined as described for the differential abundance analysis and also CLR-transformed. We calculated the correlation between the gene counts and bacterial relative abundances, then filtered for r >0.4 and Benjamani-Hochberg-adjusted p-value < 0.05. The resulting genes were manually inspected for their most relevant GO annotation and grouped according to their immunological function and pro-or anti-inflammatory nature (Supplemental table S1).

Normalized gene counts were used to perform random forest feature selection as implemented in the Boruta R package 7.0.0 (46). Only the importance measures of statistically significant (p<0.01) features were reported.

### Statistical Analyses

All statistical analyses were performed in R version 4. Alpha diversity comparisons were evaluated using the Wilcoxon rank sum test. Beta diversity was compared using Permutational Multivariate of Variance (PERMANOVA) using the adonis2 function of the vegan R package. The relationship between treatment arm and CST was assessed using multinomial logistic regression. RNAseq and 16S taxa correlations were calculated using Pearson coefficient. A significance threshold of • = 0.05 was used for the differential abundance hypothesis testing.

### Data Availability

The 16S raw reads and RNASeq libraries will be deposited in the National Center for Biotechnology Information Short Read Archive and European Bioinformatics Institute (respectively) upon publication. R code to reproduce the analysis is available at https://github.com/bmaust/CHAPS/. The completed STORMS (Strengthening The Organizing and Reporting of Microbiome Studies) checklist (47) for this project is located at https://doi.org/10.5281/zenodo.7269027.

## Results

### Clinical STI testing

No participants reported STI symptoms, and no physical exams revealed urethral discharge or other genital abnormality. None of the participants had clinical balanoposthitis or evidence of macroscopic inflammation. No GC infections were diagnosed, but NAAT for seven participants was positive for CT: five from Uganda and two from South Africa.

### Microbiome 16S sequencing

After filtering and contamination removal, 137 specimens from the 144 enrolled participants had sufficient bacterial DNA reads to proceed with analysis. The identified bacterial taxa include a variety of skin-associated Gram-positive and genital-associated anaerobic species in addition to Gram-negative enterics (**Fig. 1**). *Corynebacterium* was the most prevalent and abundant genus, appearing in 132 (97%) of specimens at median relative abundance of 34% (range: 0.14% to 98%). Anaerobic species were highly abundant, including bacteria that are commonly found in bacterial vaginosis, an inflammatory dysbiosis of the vagina, including *Prevotella, Anaerococcus, Finegoldia* and *Porphyromonas*. There were no ASVs in the *Chlamydiaceae* family which includes *C. trachomatis*.

**Fig. 1.**
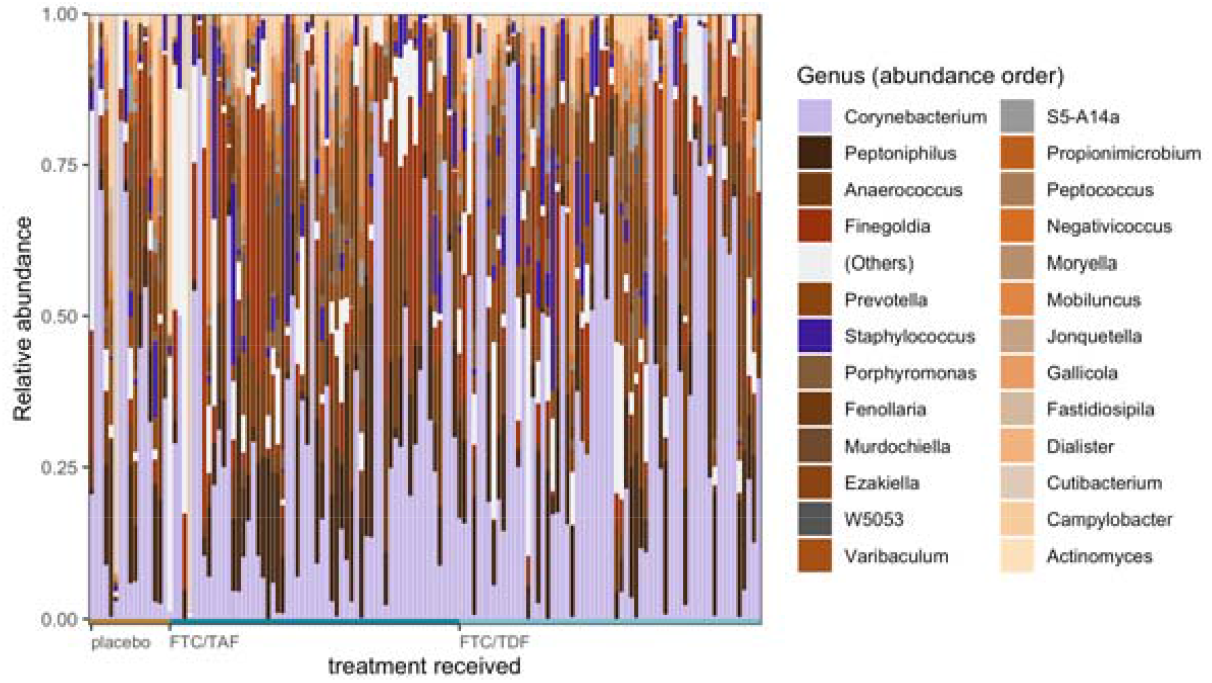
Top 25 most abundant bacterial genera across participants by study treatment received. (anaerobic taxa in earth tones, aerobes in purples, uncultured or unidentified species in grays)

Unsupervised partition around medioids clustering separated the unweighted Unifrac distances into two community structure types (CST) with distinct community structure (PERMANOVA R^2^ = 0.25 with p < 0.001) and alpha diversity (p=2.39×10 ^-19^) (**Fig. 2**). Two clusters maximized the silhouette score with acceptable within sum of squares and gap statistics. CST1 was highly diverse with a median Shannon index of 3.05. CST2 was dominated by *Corynebacterium tuberculostearicum* and *Finegoldia magna* and, with median relative abundances of 21% and 9.6%, respectively. CST2 also had a significantly lower median Shannon index of 1.68 (Wilcoxon unpaired exact, p = 2.39×10^−19^). Though *F. magna* and *C. tuberculostearicum* were also abundant in CST1 (median relative abundances of 4.6% and 2.5%), they shared high relative abundance with *Anaerococcus, Campylobacter, Fenollaria, Finegoldia, Ezakiella, Mobiluncus*, and *Peptinophilis* species without a clear dominant taxon.

**Fig. 2.**
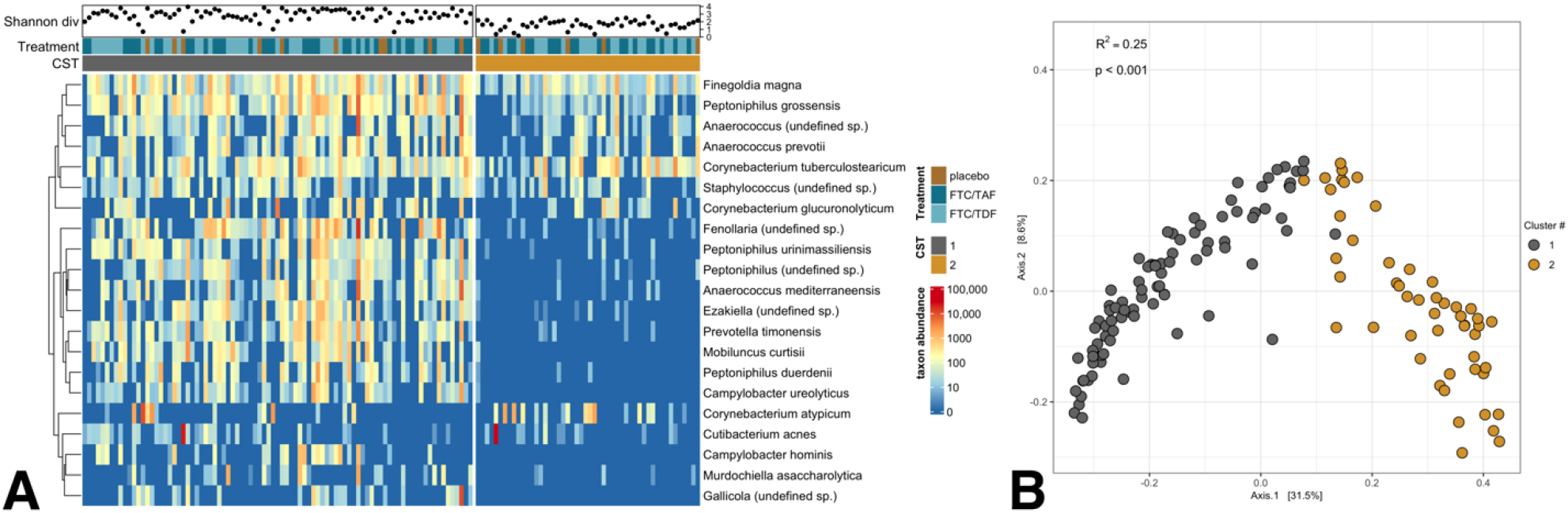
(A) Heatmap of top 25 bacterial species by CST, (B) PCoA ordination of bacterial beta diversity by unweighted Unifrac distance, colored by CST

### Microbiome differences by study site

The 137 participants with 16S data included 69 (50.3%) individuals from South Africa and 68 (49.7%) from Uganda. We compared microbiota between the two study sites and found no differences in within-participant alpha diversity (Shannon entropy) or between-participant beta diversity (unweighted Unifrac distance) (**Fig. 3, A and B**). Differential abundance testing identified *Cutibacterium acnes* as significantly higher in participants from South Africa compared to those from Uganda by ALDEx2 with a CLR difference of 3.3 between sites (Wilcoxon rank test with Benjamani-Hochberg correction p=0.0184). ANCOM-BC identified the same ASV with a significant q-value (0.0079), but the log_2_fold change of 1.87 failed to meet the effect size threshold. DESeq2 did not identify any taxa as significantly differentially abundant. (**Fig. 3, C-E**).

**Fig. 3.**
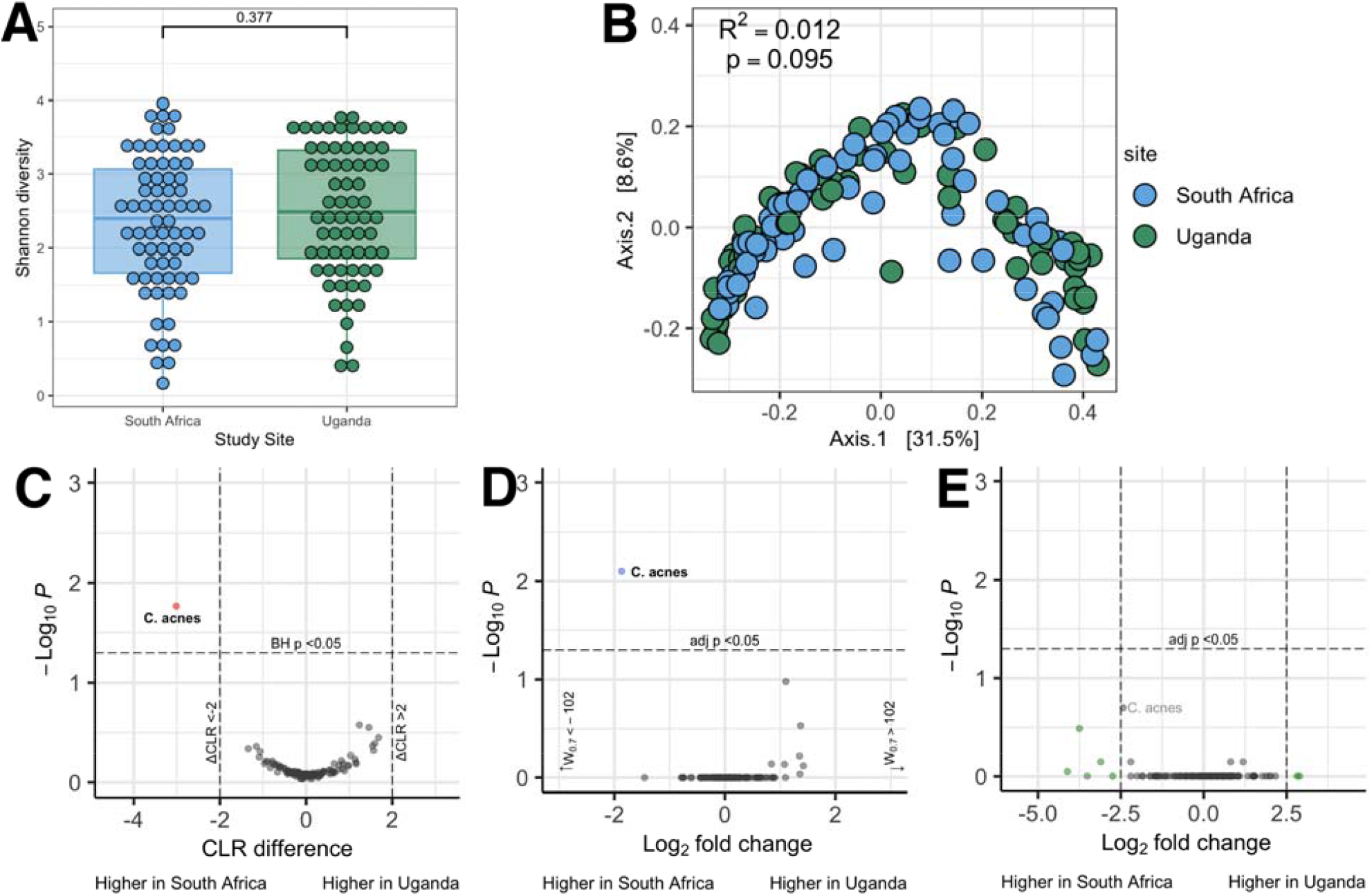
Comparisons by study site: alpha (A) and beta (B) diversity; differential abundance testing with ALDEx2 (C), ANCOM-BC (D), and DESeq2 (E)

### Microbiome differences by parent study treatment arm

We performed similar analyses for the bacterial populations in participants who received active drug versus placebo. The participants with 16S data were equally distributed among treatment arms with FTC-TAF and FTC-TDF (n=59 and 62, respectively, •^2^ p=0.437). All 16 participants who received placebo had sufficient sequences for analysis. We found no differences in alpha or beta diversity (Fig. **4**, A and B) between placebo and FTC-TAF or FTC-TDF regimens.

Combining both treatment groups and comparing to placebo, no species were significantly differentially abundant by any of the three tools (**Fig. 4, C-D**). An un-annotated *Dialister* species was identified with statistically significantly higher abundance in participants who received active drug by ANCOM-BC but did not meet the effect size threshold at only 1.9-fold more abundant. The treatment arm was not a significant predictor of the CSTs identified above (p=0.32 for placebo, p=0.99 for FTC /TAF, p=0.88 for FTC /TDF).

**Fig. 4.**
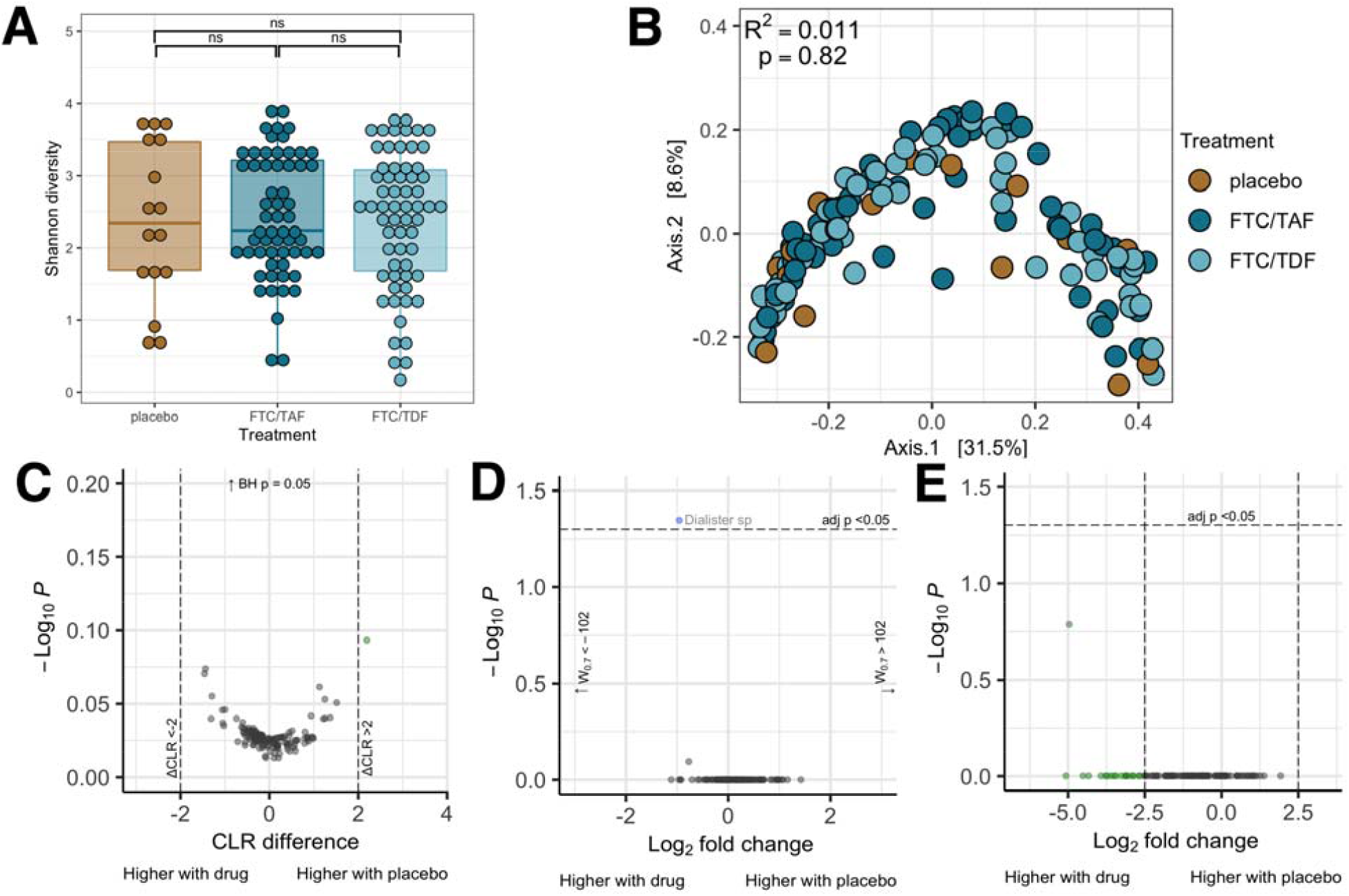
Analysis by treatment arm: alpha (A) and beta (B) diversity by drug received; differential abundance testing with ALDEx2 (C), ANCOM-BC (D), and DESeq2 (E)

### Inflammatory genes and bacterial taxa

Forty inflammatory genes showed significant correlation with 31 bacterial species (**Table 1**). Six genes had insufficient evidence for inflammatory function and were therefor excluded. The remaining 34 genes were primarily pro-inflammatory with negative correlation to bacterial species (**Fig. 5**) without difference by environmental niche. IL-15 was the most frequently correlated gene, with significant negative correlations to seven bacterial taxa not typically associated with invasive infection: *Brevibacterium luteolum, Corynebacterium urealyticum, Dietzia timorensis*, and unannotated species in the *Cutibacterium, Corynebacterium, Dietzia*, and *Nosocomiicoccus* genera. The majority of the bacterial taxa significantly correlated with other genes were gram-positive organisms which frequently colonize the skin. The CLR-transformed relative abundance of *Corynebacterium massilliense*, in particular, was associated with significantly lower expression of genes involved in regulation of inflammatory responses and neutrophil chemotaxis and activation. Anaerobes also found in the oral cavity such as *Parvimonas* and *Porphyromonas* were also correlated with primarily lower expression of inflammatory genes. However, the oral anaerobe *Rothia amarae* was associated with higher expression of the regulatory factor *GHSR* and an unclassified species also in the *Rothia* genus was associated with higher expression of the pro-inflammatory gene *REG3G*. Species in three genera canonically associated with BV, *Atopobium, Prevotella*, and *Sneathia*, were correlated with lower expression of inflammatory genes, but one unannotated *Prevotella* ASV was negatively correlated with *ZFP36*, an immune regulatory gene.

**Table 1.**
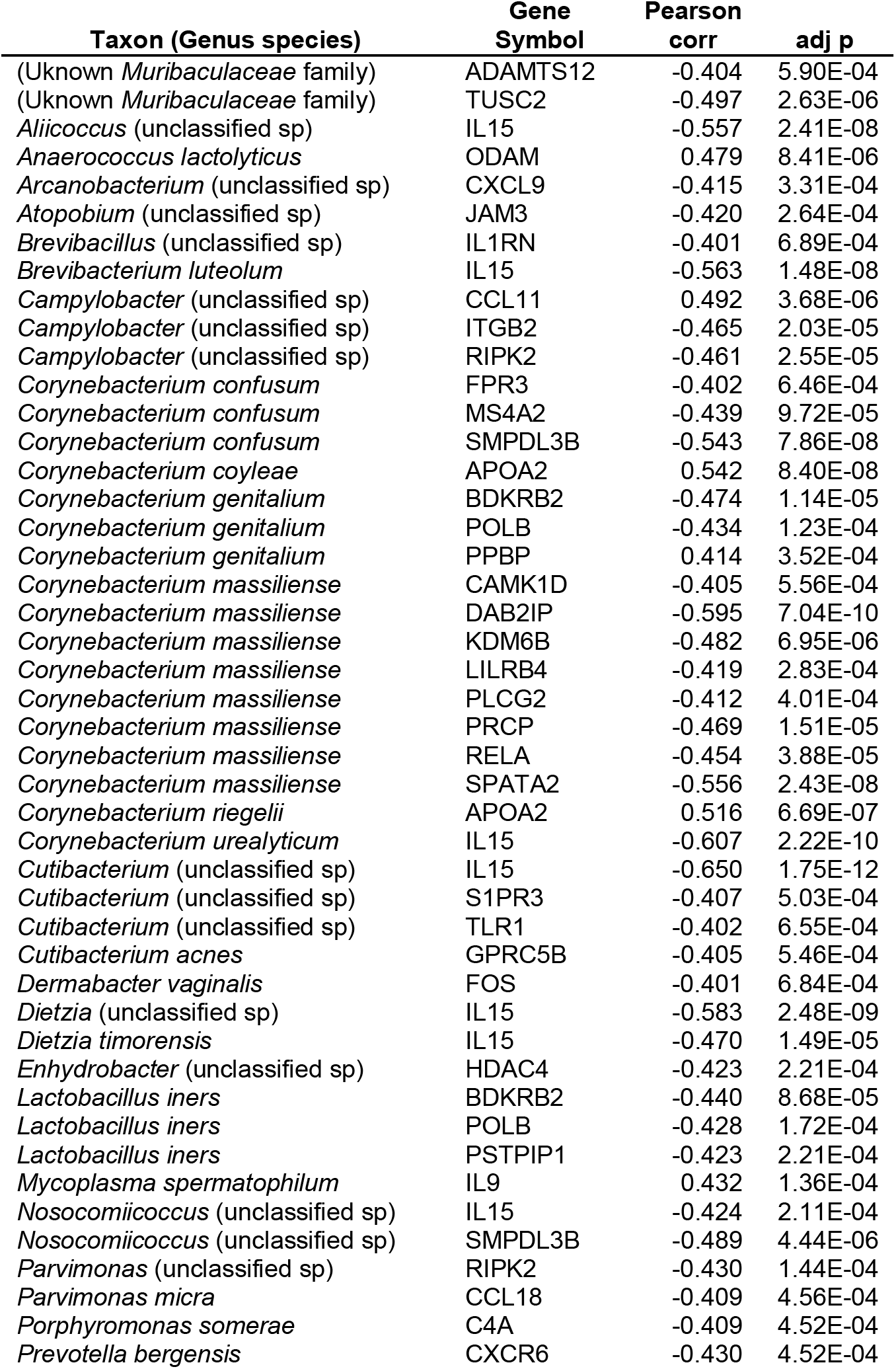
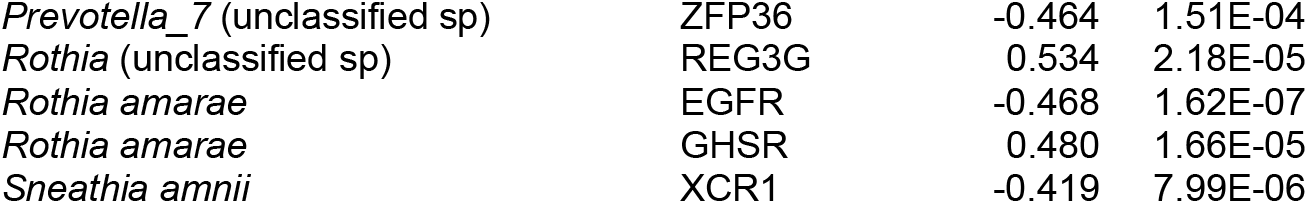
Bacterial taxa and human gene correlations

**Fig. 5.**
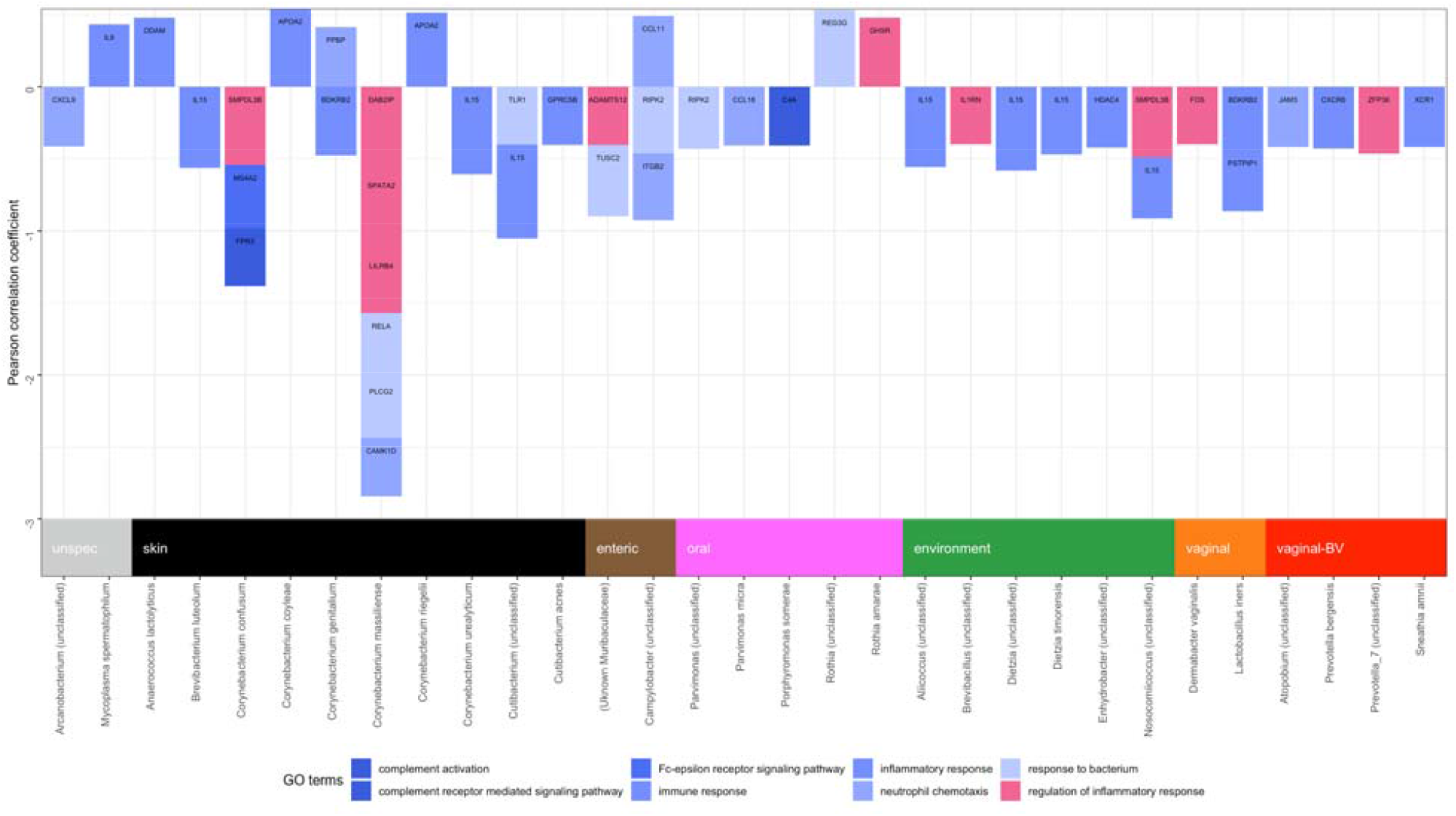
Bacterial taxa and inflammatory gene associations by immune function

We performed a similar analysis, grouping bacterial ASVs at the genus level (Supplemental table S2). Eight genes showed correlation with nine bacterial genera. As expected, all the correlated genes and bacterial genera were also identified in the species-level analysis.

We conducted a separate query of associations between inflammatory genes and the CSTs described in **Fig. 2**A distinguished by high and low diversity bacterial populations. In the random forest classification, four features achieved statistical significance. Nuclear Factor of Activated T cells 3 (*NFATC3*) and Selenoprotein S (*SELENOS*) showed higher expression in the highly diverse CST1 relative to CST2, while Signal Transducing Adapter Family Member 1 (*STAP1*) and Nod-like Receptor Pyrin domain-containing 6 (*NLRP6*) showed lower expression (**Fig. 6**).

**Fig. 6.**
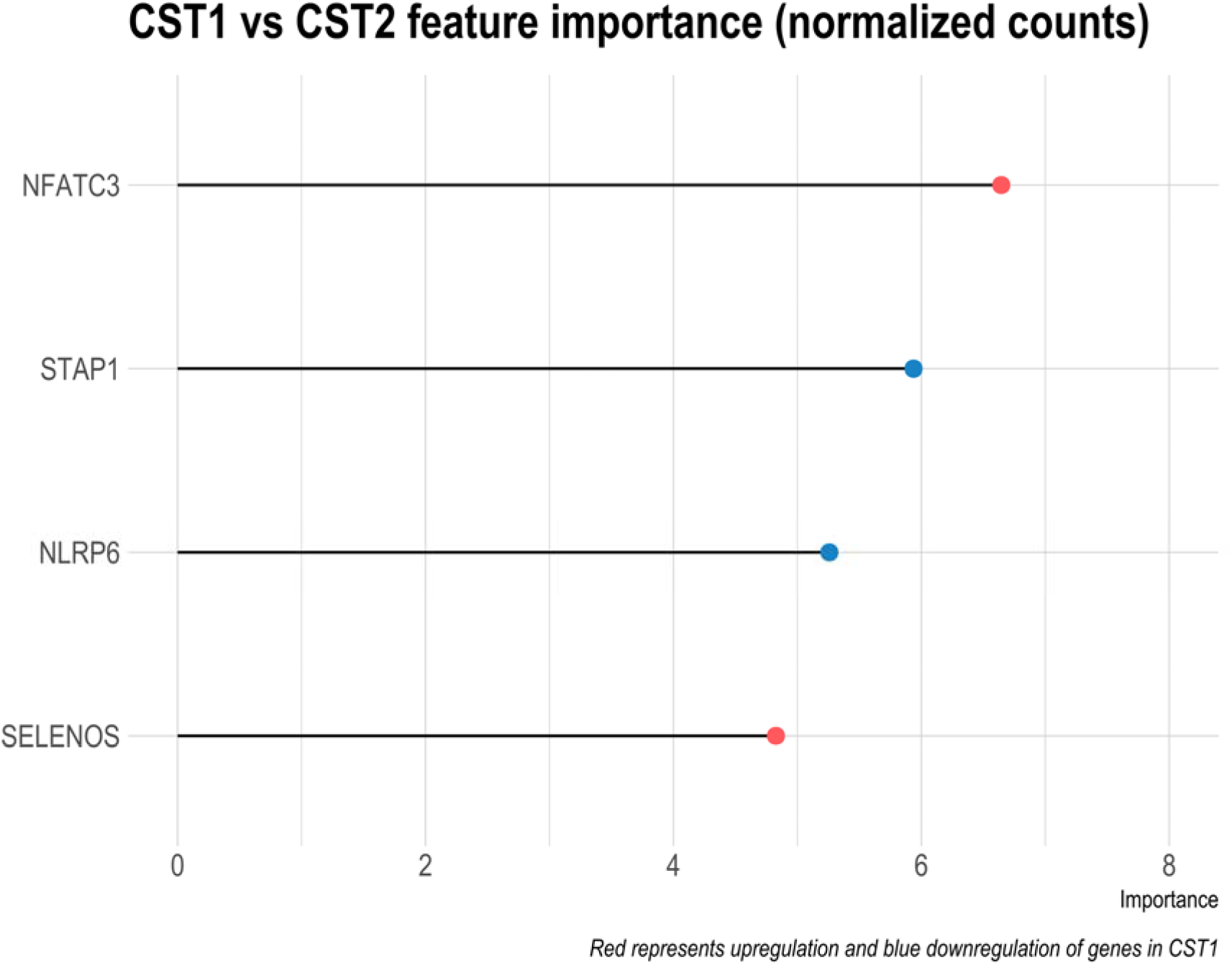
Random Forest Feature Importance comparing RNAseq results between CST 1 and CST 2

## Discussion

Our study is the first of which we are aware to analyze the tissue-level microbiome of the foreskin. Consistent with previous reports using penile swabs (5-18, 48), the bacteria we identified are predominated by taxa commonly colonizing the skin (chiefly *Corynebacteria* spp) (49, 50) in addition to anaerobic bacteria such as *Prevotella, Dialister, Murdochiella, Peptoniphilus*, and *Negativicoccus*. These anaerobic species have been associated with increased inflammation and HIV acquisition in uncircumcised men (8) and bacterial vaginosis in women (51). In the foreskin, we describe two major CSTs, one significantly more diverse than the other. We did not identify bacterial species that were differentially abundant between participants receiving placebo compared to emtricitabine with either of the two forms of tenofovir. While vaginal-associated species such as *Lactobacillus* and *Gardnerella* take up tenofovir from their environment (19, 20), further studies of more prolonged ARV use may better elucidate whether there is an effect on bacterial or viral communities of the penis.

The overall composition of the bacterial community did not appear to differ by study site; however, we did find *Cutibacterium acnes* to be significantly more abundant in South African than Ugandan participants. *C. acnes* is typically resident in the deep dermis in association with sebaceous glands and hair follicles (52), which our study sampled by digesting full-thickness specimens rather than resuspending skin swabs. While its contribution to its namesake acne vulgaris is debated, it is otherwise non-pathogenic in immunocompetent hosts without artificial material (52). It is frequently identified in surgical cultures, with unclear significance to sterility, likely due to its resistance to surgical sterilization techniques and transection of deep dermal structures during surgery (53). The differential abundance we observed may have been caused by different surgical preparation at the two sites or by an actual difference in bacterial populations. As there were no surgical complications observed during the study, the difference does not appear to have clinical significance.

Our inflammatory gene analysis primarily identified an inverse relationship between expression of genes associated with response to bacteria and skin commensals such as *Corynebacterium* and *Cutibacterium*, consistent with non-inflammatory colonization. IL-15, the most commonly correlated gene, is a pleotropic cytokine secreted by a narrow range of cell types, in foreskin tissue including epithelial cells, fibroblasts, Langerhans cells, and monocytes (54). It has broad immunostimulatory function, promoting NK cell differentiation and survival (55), inflammatory cytokine production by macrophages (56) and dendritic cells (57), neutrophil activation, survival, and phagocytosis (58), germinal center B cell proliferation (59), CD8^+^ T cell survival (60), and it is required for development of skin-resident memory CD8^+^ T cells (61). Its lower expression correlating with higher relative abundances of non-pathogenic bacteria is consistent with reduced inflammatory signaling corresponding to increased bacterial growth.

When examining gene expression between the participant groups with higher and lower diversity bacterial communities, we did not confirm previous findings in the penis (8, 10, 16) and vagina (62) that high bacterial diversity, particularly with anaerobic taxa, is associated with increased inflammation. While we frequently identified anaerobic taxa, individual species showed lower prevalence than in studies using surface swabs. This may have reduced our power to detect an inflammatory association. Another explanation could be in the structure of our experiment: rather than measuring secreted cytokines, we processed the entire tissue specimen for bulk RNAseq, which likely included many cells not directly interacting with bacteria or immune cells. Alternatively, the pre-procedure sterilization or the surgery itself may have preferentially removed inflammatory species. The sterilization or collection could also have altered host-bacterial interactions, though the rapid timeframe makes this explanation seem less likely. Lastly, the bacterial strains in our cohort may employ different metabolic or virulence strategies that render them less inflammatory. Unfortunately, our experiment was not structured to investigate this possibility.

In the vagina, high diversity communities are highly inflammatory (62) and associated with higher risk of adverse outcomes including HIV acquisition (63, 64) and preterm birth (65). The foreskin microbiome had two obvious CSTs, one highly diverse CST1 and a less diverse CST2. We hypothesized that the more diverse CST1 would be associated with higher levels of genes related to inflammation, as in the vagina. However, when comparing gene expression between CST1 versus CST2 specimens, random forest feature selection identified four genes that are not directly involved in the canonical innate antibacterial response pathways, such as TNF-_α_ and IL-6. While named for its key activity in T-Cell Receptor (TCR) signaling, NFAT family members play a broad role in cell differentiation in the immune system and beyond (66), including in B cells (67), Toll-like receptor (TLR) signaling in monocytes (68), and proliferation in perivascular tissue (69) and keratinocytes (70). Higher expression of NFATC3 in CST1 specimens may represent increased signaling from any of these cell types. SELENOS is up-regulated by cytokines such as IL-1• and TNF-• via NF-•B and in turns acts to suppress cytokine secretion in macrophages (71). Its higher expression in CST1 samples is consistent with increased suppression of chronic inflammation. STAP1 has the best evidence for signaling downstream of the B-Cell Receptor (BCR) (72), and lower expression could represent a response to sustained signaling (although this was not supported by the expected changes in other genes). NLRP6 is both an inducer and a component of the inflammasome, binds directly to the lipopolysaccharide of gram negative or lipoteichoic acid of gram positive bacterial cell membranes, and plays both a pro- and anti-inflammatory role in tissues such as liver, kidney, and intestine (73). Given these contrasting roles, the effects of its lower expression are difficult to predict with available data. None of the identified genes plays a first-line role in regulating bacterial sensing or inflammatory response.

In summary, we report on the first tissue-level examination of the bacterial microbiome of the foreskin, without apparent effect of brief antiretroviral drug exposure. Correlating bacterial species with RNAseq data revealed largely negative correlations with genes involved in the inflammatory response, consistent with maintenance of immune tolerance.

## Acknowledgements

This study is part of the EDCTP2 programme supported by the European Union (grant number RIA2016MC-1616). BSM was supported by NIH training grant T32HD007233. Research reported in this publication was supported by the National Institute of Allergy and Infectious Diseases of the National Institutes of Health under award number R21AI165144 to Heather Jaspan. The study received support by a grant from the Swedish Research Council (Francesca Chiodi; Vetenskapsrådet (VR) 2019-04596). The funders had no role in study design, data collection, and interpretation, or the decision to submit the work for publication. The authors declare that no conflict of interest exists.

## Supplemental items

Supplemental Table S1: Gene categorization

Supplemental Table S2: Bacterial genera and human gene correlations

