## Supplemental tables for "Bacterial Microbiome and Host Inflammatory Gene Expression in Foreskin Tissue"

**Supplemental Table S1**: Gene categorization

| **Gene Symbol** | **GO ID** | **GO annotation** | **Pro/anti-inflammatory** | **category** |
| --- | --- | --- | --- | --- |
| ADAMTS12 | GO:0050727 | regulation of inflammatory response | anti | regulation of inflammatory response |
| APOA2 | GO:0032757 | positive regulation of interleukin-8 production | pro | inflammatory response |
| BDKRB2 | GO:0050482 | arachidonic acid secretion | pro | inflammatory response |
| C4A | GO:0006956 | complement activation | pro | complement activation |
| CAMK1D | GO:0090023 | positive regulation of neutrophil chemotaxis | pro | neutrophil chemotaxis |
| CCL11 | GO:0030593 | neutrophil chemotaxis | pro | neutrophil chemotaxis |
| CCL18 | GO:0030593 | neutrophil chemotaxis | pro | neutrophil chemotaxis |
| CXCL9 | GO:0030593 | neutrophil chemotaxis | pro | neutrophil chemotaxis |
| CXCR6 | GO:0006955 | immune response | pro | immune response |
| DAB2IP | GO:0034144 | negative regulation of toll-like receptor 4 signaling pathway | anti | regulation of inflammatory response |
| FOS | GO:0007179 | transforming growth factor beta receptor signaling pathway | pro | regulation of inflammatory response |
| FPR3 | GO:0002430 | complement receptor mediated signaling pathway | pro | complement receptor mediated signaling pathway |
| GHSR | GO:0050728 | negative regulation of inflammatory response | anti | regulation of inflammatory response |
| GPRC5B | GO:0060907 | positive regulation of macrophage cytokine production | pro | inflammatory response |
| HDAC4 | GO:0071356 | cellular response to tumor necrosis factor | pro | inflammatory response |
| IL15 | GO:0035723 | interleukin-15-mediated signaling pathway | pro | inflammatory response |
| IL1RN | GO:2000660 | negative regulation of interleukin-1-mediated signaling pathway | anti | regulation of inflammatory response |
| IL9 | GO:0006954 | inflammatory response | pro | inflammatory response |
| ITGB2 | GO:0030593 | neutrophil chemotaxis | pro | neutrophil chemotaxis |
| JAM3 | GO:0090022 | regulation of neutrophil chemotaxis | pro | neutrophil chemotaxis |
| LILRB4 | GO:1900016 | negative regulation of cytokine production involved in inflammatory response | anti | regulation of inflammatory response |
| MS4A2 | GO:0038095 | Fc-epsilon receptor signaling pathway | pro | Fc-epsilon receptor signaling pathway |
| ODAM | GO:0006954 | inflammatory response | pro | inflammatory response |
| PLCG2 | GO:0032496 | response to lipopolysaccharide | pro | response to bacterium |
| PPBP | GO:0030593 | neutrophil chemotaxis | pro | neutrophil chemotaxis |
| PSTPIP1 | GO:0045087 | innate immune response | pro | immune response |
| REG3G | GO:0050830 | defense response to Gram-positive bacterium | pro | response to bacterium |
| RELA | GO:0009617 | response to bacterium | pro | response to bacterium |
| RIPK2 | GO:0050830 | defense response to Gram-positive bacterium | pro | response to bacterium |
| SMPDL3B | GO:0050728 | negative regulation of inflammatory response | anti | regulation of inflammatory response |
| SPATA2 | GO:0010803 | regulation of tumor necrosis factor-mediated signaling pathway | anti | regulation of inflammatory response |
| TLR1 | GO:0071221 | cellular response to bacterial lipopeptide | pro | response to bacterium |
| TUSC2 | GO:0050829 | defense response to Gram-negative bacterium | pro | response to bacterium |
| XCR1 | GO:0006955 | immune response | pro | immune response |
| ZFP36 | GO:0050728 | negative regulation of inflammatory response | anti | regulation of inflammatory response |
| S1PR3 | GO:0006954 | *inflammatory response* |  | Excluded |
| EGFR | GO:0006954 | *inflammatory response* |  | Excluded |
| KDM6B | GO:0006954 | *inflammatory response* |  | Excluded |
| PRCP | GO:0006954 | *inflammatory response* |  | Excluded |
| POLB | GO:0006954 | *inflammatory response* |  | Excluded |

**Supplemental table S2**: Bacterial genera and human gene correlations

| **Bacterial Genus** | **Gene Symbol** | **Pearson corr** | **Adj p** |
| --- | --- | --- | --- |
| (Unknown *Muribaculaceae*) | TUSC2 | -0.454 | 3.47E-05 |
| *Aliicoccus* | IL15 | -0.493 | 3.00E-06 |
| *Arcanobacterium* | CXCL9 | -0.413 | 3.26E-04 |
| *Dietzia* | IL15 | -0.463 | 2.03E-05 |
| *Lactobacillus* | BDKRB2 | -0.401 | 6.10E-04 |
| *Mycoplasma* | IL9 | 0.431 | 1.29E-04 |
| *Nosocomiicoccus* | SMPDL3B | -0.466 | 1.69E-05 |
| *Prevotella*_7 | ZFP36 | -0.441 | 7.37E-05 |
| *Rothia* | GHSR | 0.427 | 1.63E-04 |
